# Somatically Mutated Genes Under Positive and Negative Selection Found by Transcriptome Sequence Analysis Include Oncogene and Tumor Suppressor Candidates

**DOI:** 10.1101/396739

**Authors:** Casey W. Drubin, Avinash Ramu, Nicole B. Rockweiler, Donald F. Conrad

## Abstract

**Introduction:** Oncogenic somatic mutations confer proliferative advantage and undergo positive clonal selection. We developed software and applied new analytical approaches to identify: (1) somatic mutations in diverse tissues, (2) somatically mutated genes under positive and negative selection, (3) post-transcriptional modifications in the mitochondrial transcriptome, and (4) inherited germline alleles predisposing people to higher somatic mutation burden or higher levels of post-transcriptional modification.

**Methods:** Transcriptome sequence data (Genotype Tissue Expression project) for 7051 tissue samples from 549 postmortem donors and representing 44 tissue types were used. Germline mutations were inferred from whole-exome DNA sequencing and SNP arrays. DNA somatic mutations were inferred from variant allele frequencies (VAF) in RNA-seq data. Post-transcriptional modifications were inferred from Polymorphism Information Content (PIC) at the p9 sites of mitochondrial tRNA sequences. Positive and negative clonal selection was evaluated using a nonsynonomous/synonomous mutation rate (dN/dS) model. Genome-wide association studies (GWAS) were assessed with mitochondrial PIC for post-transcriptional modification level, or using the total number of somatic mutations observed per donor for somatic mutation burden.

**Results:** Our dN/dS model identified 78 genes under negative selection for somatic mutations (dN/dS < 1, p_adj_ < 0.05) and 14 under positive selection (dN/dS > 1, p_adj_ <0.05). Our GWAS identified 2 sites associated with post-transcriptional modification (1 approaching significance with p=5.99×10^−8^, 1 with p<5×10^−8^) and ∼20 sites associated with somatic mutation burden (p<5×10^−8^).

**Conclusions:** To our knowledge these are the first genome-wide association studies on normal somatic mutation burden. These studies were an attempt to increase understanding of the somatic mutation process. Our work identified somatic mutations at the global organismal level that may promote cell proliferation in a tissue-specific manner. By identifying tissue-specific mutations in actively expressed genes that appear before cancer phenotype is detected, this work also identifies gene candidates that might initiate tumorigenesis.

## Introduction

Mutations that confer or deny a proliferative advantage, including those in oncogenes and tumor suppressors, will become enriched in tissues from many organs of the body and from many individuals as a function of age. These mutations may arise from UV and oxidative damage or because of errors in replication during tissue renewal. The combined impact of environmental mutagens and tissue renewal lead to accumulation of somatic mutations as a function of age, and selection can give progenitor cells and their progeny a growth advantage.

However, still uncharacterized are many genes, including but not limited to oncogenes and tumor suppressors, which promote proliferative growth when mutated. The tissue samples investigated in this project are not from cancer tissue, but rather are from normal tissues, which accumulate somatic mutations over time.**^1^** The tissue-specific expression profile of early oncogenic mutations remains poorly understood. One approach to identifying genes that control cell proliferation and survival, some of which may be novel oncogenes and tumor suppressors, is to identify genes in which somatic mutations show hallmarks of strong positive selection. The ratio of non-synonymous to synonymous point mutations, known as the dN/dS ratio, compares the observed ratio of non-synonymous to synonymous mutations against the theoretical ratio achieved by enumerating all possible point mutations in each codon of a gene of interest.**^2^** This method has been used often in evolutionary biology and is increasingly used in cancer biology.**^3^** In the case of strong positive selection for somatic mutations, non-synonymous (missense) mutations are predicted to become proportionally overrepresented, while synonymous (silent) mutations are predicted to become proportionally underrepresented. Thus, in the case of positive selection, it is expected that most often dN/dS > 1. Similarly, in the case of negative selection, the predictions are that generally dN/dS < 1 and in the absence of selection dN/dS = 1. Here, we use dN/dS models in conjunction with genome-wide association studies (GWAS) and sequencing technology that makes it possible to obtain whole-transcriptome data for numerous tissues.**^4^** We also validate our somatic mutation call-set and our donor genotyping by analyzing genes involved in mitochondrial post-transcriptional modification.^5^

## Methods

RNA-seq data were obtained from the Genotype Tissue Expression project (GTEx V6p release). This dataset contains 550 postmortem donors with 7051 tissue samples across 44 tissue types. Germline mutations were inferred from whole-exome DNA sequencing and SNP arrays in the GTEx library. Somatic mutations were inferred from variant allele frequency (VAF) in transcriptome-wide RNA sequencing (**equation 1**: *N* is the total sample coverage, *k* is the alternative allele coverage, and *p* is the background error rate from sequencing techniques). Post-transcriptional modification was inferred from polymorphism information content (PIC) at the p9 sites of mitochondrial tRNA sequences (**equation 2**: *f* is the fraction of RNA-seq reads corresponding to a nucleotide).

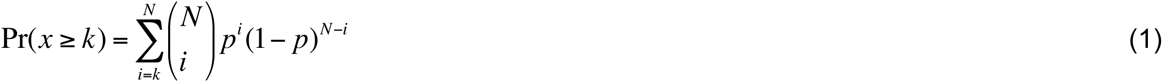

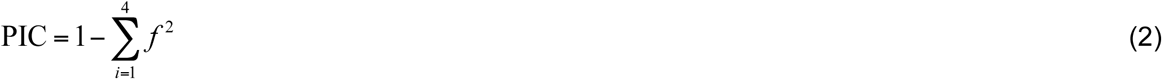

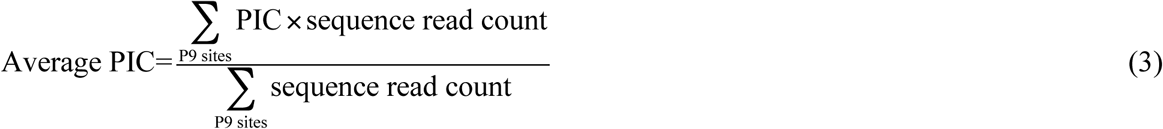

PIC was calculated for each tissue sample at each mitochondrial base pair called as a variant (**equation 1**). PIC was observed to be > 0.5 at known mitochondrial methylation positions in the 9^th^ base pair of mitochondrial tRNA genes positions (**Fig 2A**).**^5^** PIC at mitochondrial genome positions 1610, 2617, 5520, 7526, 8303, 9999, 10413, 12146, 13710, and 14734 (relative to hg19) was used (**Figs 1C & F**). The phenotype in the GWAS for RNA editing was a weighted average PIC value (**equations 2-3**). The phenotype in the GWAS for somatic mutations was the number of somatic mutations per megabase of RNA-seq, scaled to a value between 0 and 1. All GWAS was performed using PLINK v1.07.**^6,7^** Germline variants selected for the analysis had mean allele frequency (MAF) > 1%, mean allele count (MAC) > 3, and p(Hardy Weinberg Disequilibrium) < 0.1%. Covariate analysis was used to eliminate population stratification and a linear association model was used. The threshold for declaring statistical significance was 5×10^−8^, a benchmark that is widely used in the GWAS community.

**Figure 1:**
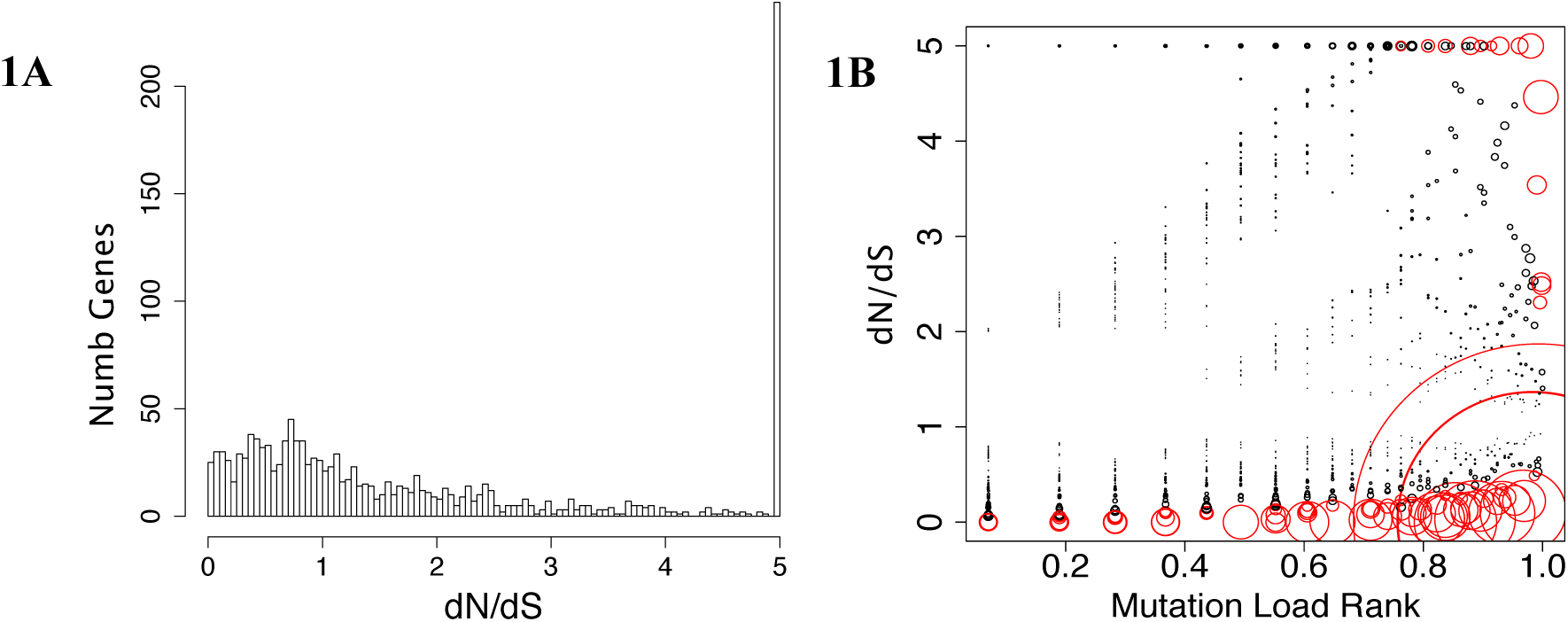
**A.** Histogram of dN/dS value for genes with ≥ 6 somatic mutations. The data appear to follow a Poisson distribution with median=1.24 **B** dN/dS value vs. mutational load rank scaled from 0 to 1 for each gene with ≥ 6 somatic mutations. The size of each circle depicts significance. Red circles are significant: p_adj_<0.05 at FDR=5%.

**Figure 2:**
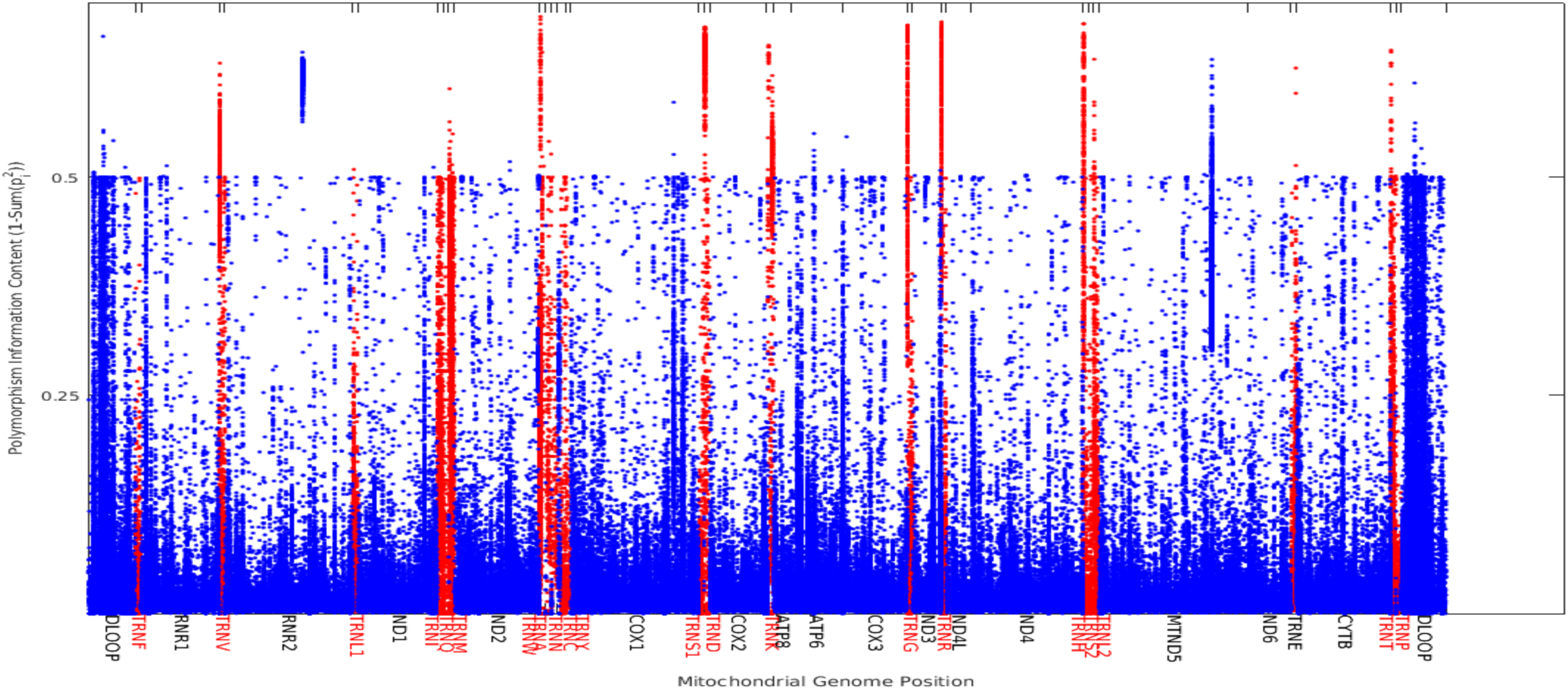
PIC value across the mitochondrial genome. tRNA positions are highlighted in red (note peaks in those positions). The 9^th^ position of each tRNA was used as the phenotype in the post-transcriptional GWAS.

While inherited alleles associated with increased incidence of somatic mutations and post-transcriptional modification were assayed using GWAS, putative somatic oncogenes were assayed using a dN/dS model (**equation 3**).

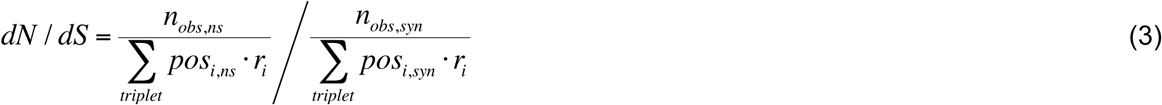

n_obs_ represents the total number of observed somatic mutations, either synonymous (silent) or nonsynonymous (missense), for a given gene across all tissue samples. In order to calculate pos_i_, the number of possible mutations falling into 1 of 96 classes, every SNP was simulated at every position of any gene with observed mutations. The 96 classes of SNPs represent 3 possible nucleotide changes in all 64 codons, but dividing by the 2-fold degeneracy of dsDNA. Since different classes of mutations happen with different likelihood (for example transition vs. transversion), rate constants r_i_ were used to adjust the potential-mutation counts and generate expected-mutation counts. Rate constants were obtained for each of the 96 mutation classes by summing all observed mutations and dividing by the sum of all possible mutations across all genes sequenced. Annovar (v2016Feb01) was used to assign a gene ID to all mutations and annotate them as synonymous or nonsynonymous.**^8^** P-values were obtained for all dN/dS values using Poisson regression. Adjusted p-values were obtained using a false discovery rate (FDR) of 5% and significance threshold of p adj<0.05. Only genes with ≥ 6 somatic mutations (n=1385 genes) were used in the analysis of dN/dS as genes with fewer mutations tended not to rise to statistical significance. Muscle and sun-exposed skin were not used in the dN/dS analysis as these tissues were observed to have different mutation rate constants r_i_ from the majority of tissues.

## Results

### dN/dS Model

Our work to identify genes that might initiate oncogenesis used a dN/dS model to investigate transcriptome data at the tissue-specific level. **Table 1** summarizes all genes found to be significant for positive selection. This table also highlights the tissues in which genes are expressed, one of the important features of our dataset. As shown in **Table 1**, the 2 genes with dNdS most significantly > 1 (lowest p value) are ARRB2, a beta arrestin gene, and CSF3R, a leukemia-associated colony stimulating factor gene. The ARRB2 mutations were found exclusively in whole blood (ARRB2 is expressed most highly in whole blood, but is also expressed in numerous other tissues such as adipose, brain, and spleen) and were observed in 50 of the 549 donors. This gene is important in the Wnt signaling pathway and is an oncogene.**^10^** Similarly, CSF3R is implicated in leukemia.**^11^** Mutations were observed in lung and whole blood tissues from 104 individuals. Interestingly, lung tissue is typically contaminated with substantial amounts of blood, so the enrichment of CSF3R mutations seen in lung may be ultimately attributable to biological processes happening in blood. Twelve other genes with dNdS>1 reached significance (p_adj_<0.05) using FDR=0.05. For example: CARD19 (C9orf89), a Caspase recruitment protein (mutations observed in 23 donors in whole blood and lung samples; p_adj_=0.0048); LZTS2 (leucine zipper tumor suppressor 2, observed in nerve and artery tissues and in 20 different donors, p_adj_=0.017); and SPI1 proto-oncogene (observed in 26 different donors in lung and whole blood p_adj_=0.049) all reached significance. As shown in Table 1, several of these genes have known involvement in cancer. Furthermore, most genes in Table 1 have dNdS > 5, showing that the observed proportion of nonsynonymous mutations is more than five times greater than would be expected if somatic mutations were not under positive selection.

**Table 1:** dN/dS results are summarized for 14 significant (FDR 5%) genes with dN/dS>1. The top 2 genes in the table met the Bonferroni correction factor (p<0.05, n=1385). *Not depicted*: 78 genes had dN/dS significantly lower than 1 (FDR 5%), 34 of which met Bonferroni correction.

| Gene | Function (Ref 9) | dN/dS | S. Obs | NS. Obs | S. Pos | NS. Pos | p<sub>adj</sub> | Tissues | # Donors |
| --- | --- | --- | --- | --- | --- | --- | --- | --- | --- |
| ARRB2 | Beta-arrestin (signaling) | 9.41 | 2 | 64 | 989.7 | 3366.6 | 5.20e-4 | blood | 50 |
| CSF3R | Colony stimulating factor | 4.46 | 8 | 124 | 1852.1 | 6432.5 | 3.73e-5 | blood & lung | 104 |
| CARD19 (C9orf89) | Caspase recruitment protein. Inhibits NFkB | — | 0 | 28 | 494.85 | 1313.2 | 0.0048 | blood & lung | 23 |
| LZTS2 | Leucine zipper tumor suppressor | — | 0 | 23 | 1769.7 | 4669.5 | 0.0167 | Nerve & artery | 20 |
| SPI1 | Proto-oncogene. Myeloid & lymphoid development | 9.73 | 1 | 32 | 804.00 | 2643.1 | 0.0495 | Lung & whole blood | 26 |
| BDH2 | 3-hydroxybutyrate dehydrogenase | — | 0 | 18 | 659.22 | 1670.7 | 0.0499 | Many, whole body | 18 |
| SPARC | Proteolytic matrix remodeling; tumor invasion | 7.58 | 2 | 45 | 737.12 | 2186.6 | 0.0072 | Many, whole body | 40 |
| SNX3 | Nexin family intracellular trafficking | 10.4 | 1 | 30 | 482.63 | 1390.4 | 0.0303 | Heart, lung, blood, lymphocyte | 30 |
| ZYX | Focal adhesion and signal transduction | 3.54 | 8 | 77 | 1495.7 | 4067.6 | 0.0036 | Many, whole body | 66 |
| TG | Thyroglobulin | 2.52 | 17 | 120 | 6965.2 | 19512 | 0.0033 | Thyroid, adrenal, brain, esoph, fibroblast | 34 |
| RPL12 | Ribosomal protein | — | 0 | 21 | 479.72 | 1138.3 | 0.0218 | Many, whole body | 26 |
| A2M | Protein cleavage | 2.31 | 16 | 98 | 3843.9 | 10209 | 0.0202 | Many, whole body | 62 |
| NDE1 | Neurodevelopment protein, centrosome and dynein interaction | 2.48 | 17 | 124 | 5837.5 | 17141 | 0.0042 | Artery & colon | 3 |
| CSDC2 | Binds histone mRNA | 13.9 | 1 | 36 | 427.74 | 1107.8 | 0.0042 | Adipose, artery, heart, ovary, uterus | 28 |

**Fig 1A** shows the distribution of dN/dS for all genes studied across all tissues. The majority of genes appear to be under neutral or slightly positive selection for somatic SNPs. Median dN/dS = 1.2 for genes with ≥ 6 somatic mutations and the data appear to follow a Poisson distribution. However, dN/dS can only be assessed with mutations that are at a high enough tissue-wide frequency to detect *and* that are in sufficiently expressed genes. These widely distributed and expressed mutations are candidates to be important for oncogenesis. While the majority of mutations appear to be under fairly neutral selection, **Fig 1B** shows that some mutations are under positive or negative selection, with more significant results in the category of negative selection.

### Somatic Mutations Validated and Genes Responsible for Mitochondrial Post-Transcriptional Modification Discovered

In order to validate our somatic mutations call set (**equation 1** using GTEX v6p), to examine whether our call set included signals from RNA editing rather than from somatic mutation, and to use our broad dataset to uncover new information about post-transcriptional modification, we performed GWAS on mitochondrial tRNA mutations. GWAS hits (mitochondrial PIC values phenotype × somatic calls genotype) are presented in rows 1-2 of **Table 2**. The phenotype used in mitochondrial GWAS is shown in **Fig 2**, where we successfully replicated the finding that the p9 position of mitochondrial tRNA is methylated.**^5^** Critically, a tRNA methyltransferase variant (exon 1 of TRMT61B, p=1.117×10^−8^) was the most significant somatic mutation in our GWAS to explain variation in post-transcriptional modification (**Table 2**, row 1). This same gene was also found in the study we replicated and is directly mechanistically involved in tRNA modification. One of our top hits (which failed to pass the significance threshold, p = p=6×10-8) is interesting because it is a novel finding of 2 neighboring mutations in caspase genes (p=6×10^−8^), and was not reported in the original study. The caspase genes are unique in that they appear *negatively* correlated with tRNA methylation (negative β-value). Furthermore, the mutation calls we used for the tRNA study were only seen in an early, unfiltered version of our data, suggesting RNA editing was successfully filtered out from later somatic mutation data used for dN/dS and other GWAS.

**Table 2:** Summary of GWAS results. All significant positions shown, except “somatic vars×SNP array” where only top results are depicted & the 2nd “PIC value” result approaches significance.

| GWAS study | Location | Function (Ref 9) | β-value<br>(effect size) | p-value | Gene |
| --- | --- | --- | --- | --- | --- |
| **PIC Value ×**<br>**Exome Calls** | Chr2:29092850 | tRNA methylase | 0.0413 | 1.12e-8 | TRMT61B Exon 1 |
|  | Chr11:104,897,536 | Apoptosis-related caspase | -0.1673 | 5.99e-8 | CASP1 intron |
| **Somatic Vars ×**<br>**Exome Calls** | Chr1:144,989,914 | Neuroblastoma Breakpoint Family | 0.3060 | 2.44e-8 | NBPF9 intron |
|  | Chr11:93170919 | Coiled Coil Domain Containing 67,<br>promotes centriole amplification | 0.2142 | 4.81e-10 | CCDC67 Exon 14 |
|  | Chr20:18,446,026 | Double zinc ribbon and ankyrin repeat<br>domains 1 | 0.2234 | 3.32e-8 | DZANK1 intron |
| **Somatic Vars ×**<br>**SNP Array** | Chr13:91609870 | (Likely regulatory) | 0.1601 | 2.52e-10 | (Non-coding) |
|  | Chr20:18368305 | Double zinc ribbon and ankyrin repeat<br>domains 1 | 0.2297 | 2.88e-10 | DZANK1 intron |
|  | Chr10:29165089 | Uncharacterized protein | 0.2295 | 3.06e-10 | C10orf126 intron |
|  | Chr7:106403560 | unknown | 0.2460 | 5.37e-10 | AF086203 |
|  | Chr2:38409838 | (Likely regulatory) | 0.2239 | 6.96e-10 | (Non-coding) |
|  | Chr3:186747485 | Glycosyltransferase | 0.2666 | 1.25e-9 | ST6GAL1 intron |
|  | Chr2:38407303 | Oxidation and electron transport | 0.212 | 1.82e-9 | CYP1B1 intron |
|  | Chr6:114473296 | Heparan sulfate sulfotransferase | 0.198 | 2.10e-9 | HS3ST5 intron |
|  | Chr7:146723583 | Neurexin cell adhesion molecule | 0.227 | 5.06e-9 | CNTNAP2 intron |
|  | Chr16:130781408 | DNA interacting protein, interacts with<br>RB1 tumor suppressor | 0.242 | 5.71e-9 | RNF40 intron |
|  | Chr3:60846952 | Regulates cell proliferation, induce<br>apoptosis via AKT1 and SRC | 0.146 | 5.90e-9 | FHIT intron |

### GWAS Results Identify Potential Somatic Mutation-Causing Alleles

We performed 3 GWAS. We looked for (1) somatic mutation burden phenotype × exome calls genotype (QQ plot **Fig 3A**, Manhattan plot **Fig 3D**, raw genotype vs. phenotype scatter **Fig 3H**); (2) for somatic mutation burden phenotype × SNP array genotype (QQ plot **Fig 3B**, Manhattan plot **Fig 3E**); and (3) for post-transcriptional modification phenotype × exome calls genotype (QQ plot **Fig 3C**, Manhattan plot **Fig 3F**, raw genotype vs. phenotype scatter **Fig 3G**). Across QQ plots, the fact that our p-values closely track a random distribution of p-values for low and mid level p-values, but start to deviate toward higher significance for larger p-values, suggests that the studies were well calibrated. Likewise, our Manhattan plots show an acceptable level of random background association with several alleles rising to significance (p<5×10^−8^). Looking more closely at individual SNPs identified, our GWAS for variation in somatic mutation load had several novel results (**Fig 3 A,B,D,E,H** and **Table 2,** rows 3-16). A total of 3 significant alleles were identified from exome sequencing calls and a total of 59 significant alleles were identified from SNP array data. The SNP array data contains many more loci, which may explain why more significant results were found. In exome sequencing calls, the most significant hit was in a known tumor suppressor gene, exon 14 of CCDC67. This gene encodes a protein that is implicated in papillary thyroid carcinoma.**^12^** Additionally, GWAS results using exome SNPs, the 2^nd^ most significant allele was in an NBPF9 (neuroblastoma family protein 9) intron; a gene that may be associated with neuroblastoma.^13^ This NBPF9 variant was also the variant with the largest effect size (β>0.3) in the GWAS. In the SNP array, the most significant result was in a non-coding, putative regulatory region of chromosome 13. The 2^nd^ and 3^rd^ highest hits clustered together in the SNP array and were both in DZANK1, an ankyrin-coding gene on chromosome 20. Interestingly, the same gene is the third most significant gene in the GWAS performed on somatic germline mutation calls (**Table 2**), suggesting a novel role for this gene in facilitating SNPs in tissues throughout the body. Several other significant SNP array positions appeared to be cancerrelated. For example, an intronic mutation was observed in ST6GAL1, an oncogene activated by Ras, which upregulates cell migration.**^14^**

**Figure 3:**
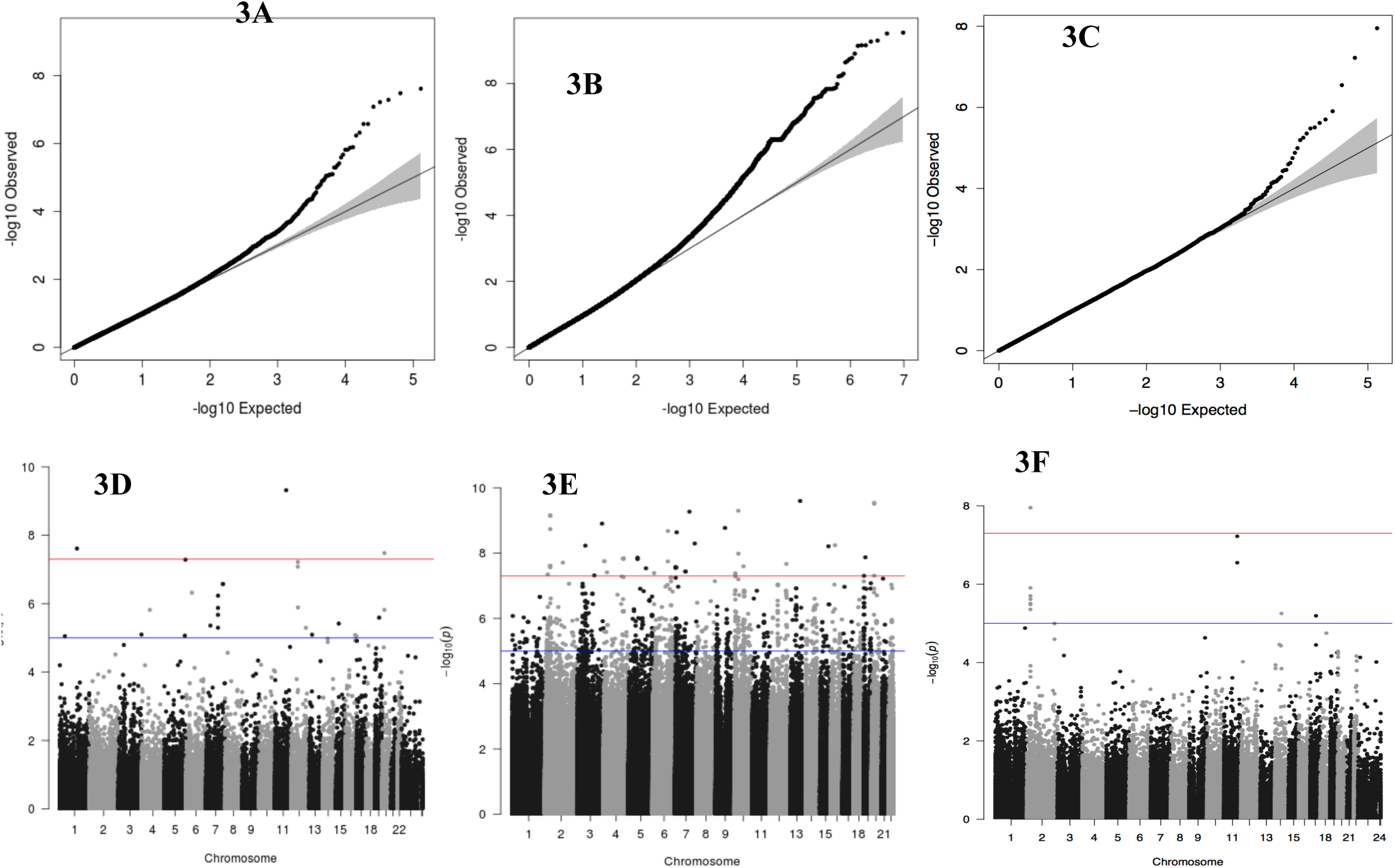
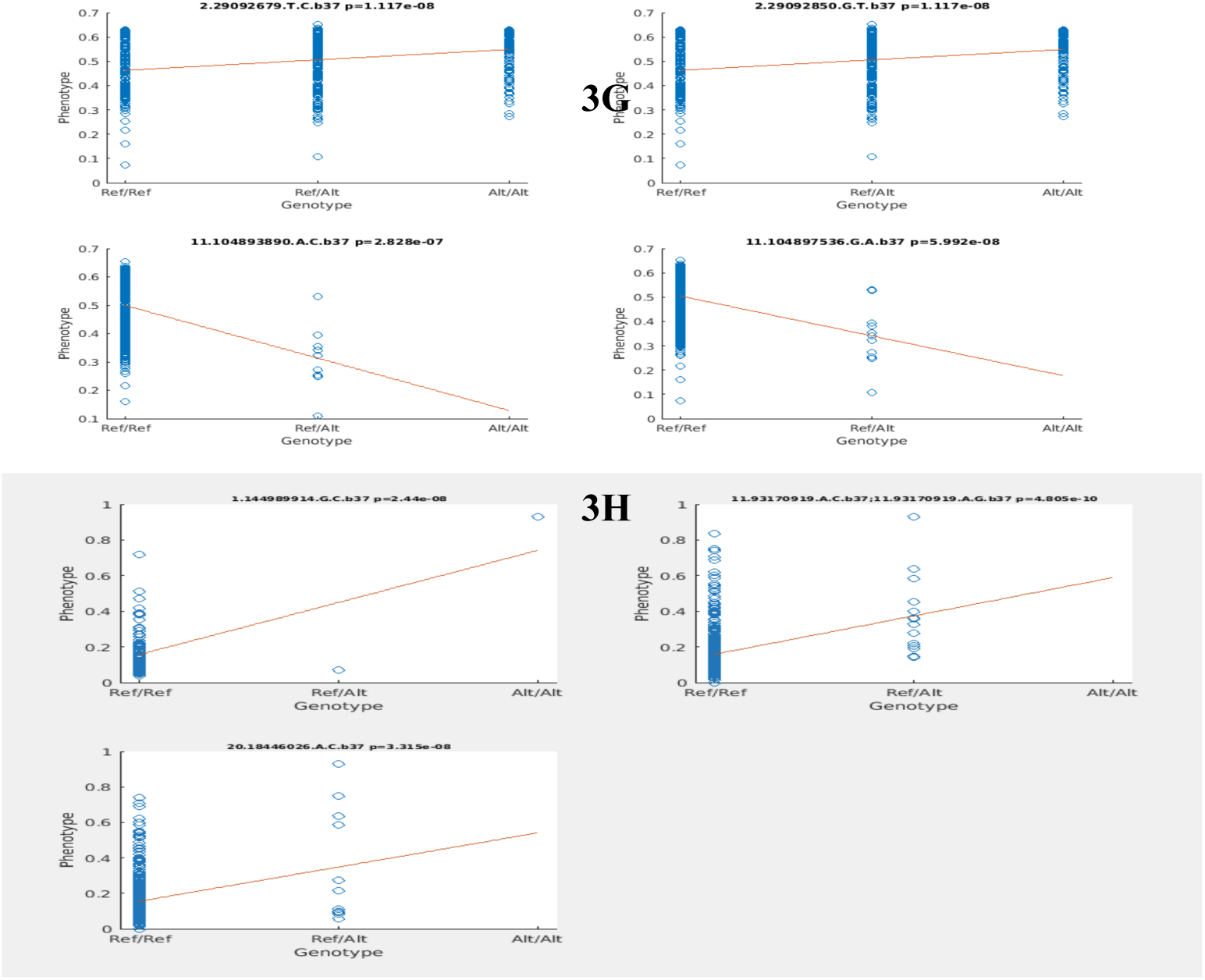
**A,B,C.** Quantile-quantile (QQ) plots for the exome mutation burden phenotype × germline variants called from seq, exome mutation burden phenotype × germline variants called from SNP array variants, and post-transcriptional modification phenotype (**equations 2-3**) × exome mutation burden respectively. **D,E,F** Manhattan plots for the same three GWAS, respectively. **G** and **H** Genotype (Ref/Ref, Ref/Alt, Alt/Alt) compared to phenotype (see *Methods*) for significant SNPs of the post-transcriptional × exome mutation burden GWAS and somatic × exome mutation burden GWAS, respectively.

## Discussion

Results presented here are consistent with the notion that tumorigenesis might be initiated differently in different tissues of the body (**Table 1**, 2^nd^ to last column). The results also identify candidates for new tumor suppressors and oncogenes (**Tables 1 and 2**). The fact that two known oncogenes were the top results for our dN/dS model reinforces the notion that oncogenes undergo positive clonal selection even in normal tissue. The median dN/dS of 1.2 across all tissues suggests that unlike Darwinian selection, which shows strong purifying (negative) selection over millions to billions of year, the forces of natural selection do not appear to act strongly on tissue mutations acquired during a person’s lifetime.**^2^** The tissue mutations observed in this study may even be under slightly positive selection since our whole-transcriptome data only includes those SNPs that are present in a high enough proportion of tissue samples *and* that are sufficiently expressed to be detected.

It is possible that some of these highly expressed SNPs are involved in initiating oncogenesis. We validated our data by replicating and adding new results to GWAS of individual variation in mitochondrial tRNA modification. The fact that we could observe mitochondrial tRNA modification and that we could find established variants suggests that our data are appropriate for answering our questions. Using GWAS, we investigated tRNA modification, but also looked into organismal-level variants that contribute to somatic mutation. Discovering these new oncogenes and novel mechanisms of tumorigenesis could open the door to new therapeutic targets for the treatment cancer and to better differentiation between healthy and diseased tissue when considering treatment.

## Acknowledgments

We thank all members of the Conrad lab for help with this project. CWD was supported by the WUSM Summer Research Program. This work was funded by grants from the NIH (www.nih.gov) [R01HD078641 and R01MH101810 to D.F.C.]

## References

1) Martincorena, I. et al. (2015) High Burden and Pervasive Positive Selection of Somatic Mutations in Normal Human Skin. Science 348, 880–886.

2) Kryazhimskiy, S. and Plotkin, J. B. (2008) The Population Genetics of dN/dS. PLOS Genet 4.

3) Ostrow, S. L. et al. (2005) Cancer Evolution Is Associated with Pervasive Positive Selection on Globally Expressed Genes. PLOS Genet 10.

4) The GTEX Consortium. (2013) The Genotype-Tissue Expression (GTEx) Project. Nat. Genet. 45, 580–5.

5) Hodgkinson, A. et al. (2014) High-Resolution Genomic Analysis of Human Mitochondrial RNA Sequence Variation. Science 344, 413–5.

6) Purcell, S. et al. (2017) PLINK v1.9. http://pngu.mgh.harvard.edu/purcell/plink/. Purcell Laboratory. Accessed 8/14/17.

7) Purell, S. et al. (2007) PLINK: A Toolset for Whole-Genome Association and Population-Based Linkage Analysis. Am. J. Hum. Genet. 81

8) Wang, K., Li, M. and Hakonarson, H. (2010). AnnOVAR: Functional Annotation of Genetic Variants from High-Throughput Sequencing Data. Nucleic Acids Res 38, e146.

9) Weizmann Institute of Science & LifeMap Sciences. (2014) GeneCards Human Gene Database. http://www.genecards.org/. Gene Cards Suite. Accessed 8/14/17

10) Bonnans, C. et al. (2012) Essential requirement for β-arrestin2 in mouse intestinal tumors with elevated Wnt signaling. Proc Natl Acad Sci USA 109, 3047–52.

11) Maxson, J.E. et al. (2013) Oncogenic *CSF3R* Mutations in Chronic Neutrophilic Leukemia and Atypical CML. N Engl J Med 368, 1781:90.

12) Yin, D.T. et al. (2016) Characterization of the Novel Tumor-Suppressor Gene *CCDC67* in Papillary Thyroid Carcinoma. Oncotarget 7, 5830–41.

13) Vandepoele, K. et al. (2005) A Novel Gene Family NBPF: Intricate Structure Generate by Gene Duplications During Primate Evolution. Mol Biol Evol 22, 2265–74.

14) Isaji, T. et al. (2014) An Oncogenic Protein Golgi Phosphoprotein 3 Up-regulates Cell Migration via Sialylation. J. Biol. Chem 289, 20694–705.

